# Magnetoencephalography Reveals Neuroprotective Effects of COVID-19 Vaccination in Non-Human Primates

**DOI:** 10.1101/2025.02.14.638187

**Authors:** Jennifer R. Stapleton-Kotloski, Jared A. Rowland, April T. Davenport, Phillip M. Epperly, Maria Blevins, Dwayne W. Godwin, Daniel F. Ewing, Zhaodong Liang, Appavu K. Sundaram, Nikolai Petrovsky, Kevin R. Porter, Christopher S. Gamble, John W. Sanders, James B. Daunais

## Abstract

COVID-19, caused by the SARS-CoV-2 virus, can lead to widespread neurological complications, including cognitive deficits and neurodegenerative symptoms, even in the absence of significant structural brain abnormalities. The potential neuroprotective effects of SARS-CoV-2 vaccination remain underexplored. Here, we demonstrate the neuroprotective effects of a psoralen-inactivated SARS-CoV-2 vaccine in a non-human primate model using resting-state magnetoencephalography (MEG), a non-invasive neurophysiological recording technique with sub-millisecond temporal and submillimeter spatial resolution. MEG scans demonstrated substantial preservation of neural activity across multiple brain regions in vaccinated subjects compared to unvaccinated controls following viral challenge. This approach not only underscores the role of vaccination in mitigating severe neurological outcomes but also highlights the capability of MEG to detect subtle yet significant changes in brain function that may be overlooked by other imaging modalities. These findings advance our understanding of vaccine-induced neuroprotection and establish MEG as a powerful tool for monitoring brain function in the context of viral infections.

## INTRODUCTION

Severe acute respiratory syndrome coronavirus 2 (SARS-CoV-2), the causative agent of COVID-19, has led to over 775 million infections and more than 7 million deaths globally as of July 2024^1^. Beyond the acute phase, a significant proportion of individuals experience persistent symptoms, collectively termed “long COVID,” which include fatigue, cognitive deficits^2–5^, and neuropsychiatric disorders such as depression, PTSD, and anxiety^6–9^. These symptoms can last for months^10–12^. Viral-specific protein has been observed in postmortem human brain tissues^13^, raising concerns over potential neuropathological consequences. Emerging evidence suggests that SARS-CoV-2 may exacerbate or even initiate neurodegenerative processes^14^, with many patients displaying marked neurological deficits^2–6^, even in the absence of detectable structural abnormalities on conventional neuroimaging^15^. Several other studies have demonstrated findings suggesting that abnormal imaging after COVID infection correlates with symptomatology. For instance, thalamic and basal ganglia changes were linked to fatigue and memory issues^16^, while grey matter atrophy corresponded to cognitive decline^14^ and vascular lesions to acute neurological events^17,18^. A multiple regression analysis of over 112,000 participants demonstrated cognitive deficits following infection with COVID-19 compared to uninfected individuals or those with unconfirmed infection, with larger deficits observed in patients with unresolved persistent symptoms and in those infected during the initial period of the COVID-19 pandemic when the wild-type Washington strain was the predominantly infectious strain^19^. Electroencephalogram (EEG) studies have revealed generalized slowing^20^ and increases in theta and alpha frequencies in those experiencing cognitive issues such as brain fog^21^.

Magnetoencephalography (MEG) is an alternative and noninvasive, clinically-used neurophysiological technique that records the biomagnetic fields generated by neural activity in real time^22^. MEG confers advantages over other modalities because unlike functional MRI (fMRI) and Positron Emission Tomography (PET), it is a direct measure of brain activity with sub-millisecond temporal resolution, and unlike EEG, possesses submillimeter spatial resolution^23^. Through the use of magnetic source imaging methods (MSI) such as synthetic aperture magnetometry (SAM), biomagnetic activity can be directly mapped in brain space^24,25^. MSI can also construct virtual electrodes (source series) for any voxel in the brain, providing full bandwidth and continuous time series of activity that correspond to the local field potentials recorded with actual invasive electrodes, enabling the detection of subtle yet clinically significant changes in neural activity.

In this study, we aimed to investigate the feasibility of using MEG to identify the neuroanatomical substrates underlying the cognitive deficits reported by COVID-19 patients, particularly in the context of post-acute sequelae. Additionally, we sought to determine whether vaccination against SARS-CoV-2 could confer neuroprotection against COVID-19-related alterations in brain function. Using a non-human primate model, we employed MEG to assess resting-state brain activity following vaccination with a novel psoralen-inactivated SARS-CoV-2 vaccine (PsIV) or the PsIV in combination with a DNA vaccine, and subsequent challenge with the Delta variant (SARS-CoV-2 B.1.617.2). Our findings indicate that vaccination significantly mitigates COVID-19-related reductions in global resting-state brain function as measured by MEG. This suggests not only a critical neuroprotective effect of vaccination but also underscores MEG’s potential as a powerful tool for detecting and monitoring the neurological impacts of infection, offering insights that conventional imaging techniques may overlook.

## RESULTS

### Immunogenicity of SARS-CoV-2 PsIV in non-human primates

In this follow-on, cross-sectional study, we leveraged NHPs that were subjects in an ongoing vaccine dose escalation study^26^ to determine feasibility of applying MEG to record resting-state (RS) brain function in female cynomolgus monkeys following vaccination with a novel psoralen-inactivated vaccine (PsIV) or a combination of a DNA vaccine and PsIV against SARS-CoV-2. The subjects were subsequently challenged with the live Delta strain (SARS-CoV-2 B.1.617.2).

Vaccination was administered in a dose-escalation study, with immunologic and virologic responses correlating with the administered dose^26^. These responses were most pronounced when the vaccine was delivered as a prime-boost regimen following DNA vaccination (Figure 1). Specifically, four groups of cynomolgus macaques (*Macaca fascicularis*), each consisting of eight animals, were immunized with adjuvant in PBS (control group) or varying titers of SARS-CoV-2 PsIV (0.075-3.75 µg) formulated with the Advax-2 adjuvant on days 0 and 30 as shown in Table 1. A fifth group of 8 animals (heterologous Prime-boost group), was vaccinated with two doses of a DNA vaccine encoding the full-length SARS-CoV-2 spike protein on days -30 and 0, followed by a booster dose of SARS-CoV-2 PsIV (3.75 µg) on day 30. Blood samples were collected on days -30 (only from the prime-boost group), 0, 30, 35, and 51, and serum was tested for the presence of anti-SARS-CoV-2 neutralizing antibodies using microneutralization assays against both the Washington strain and the SARS-CoV-2 Delta variant. The psoralen-inactivated SARS-CoV-2 vaccine with Advax-CpG adjuvant demonstrated strong, dose-dependent neutralizing antibody responses against both the Washington strain and Delta variants in nonhuman primates, with the highest protective efficacy seen when used as a booster following DNA vaccination^26^.

**Figure 1:**
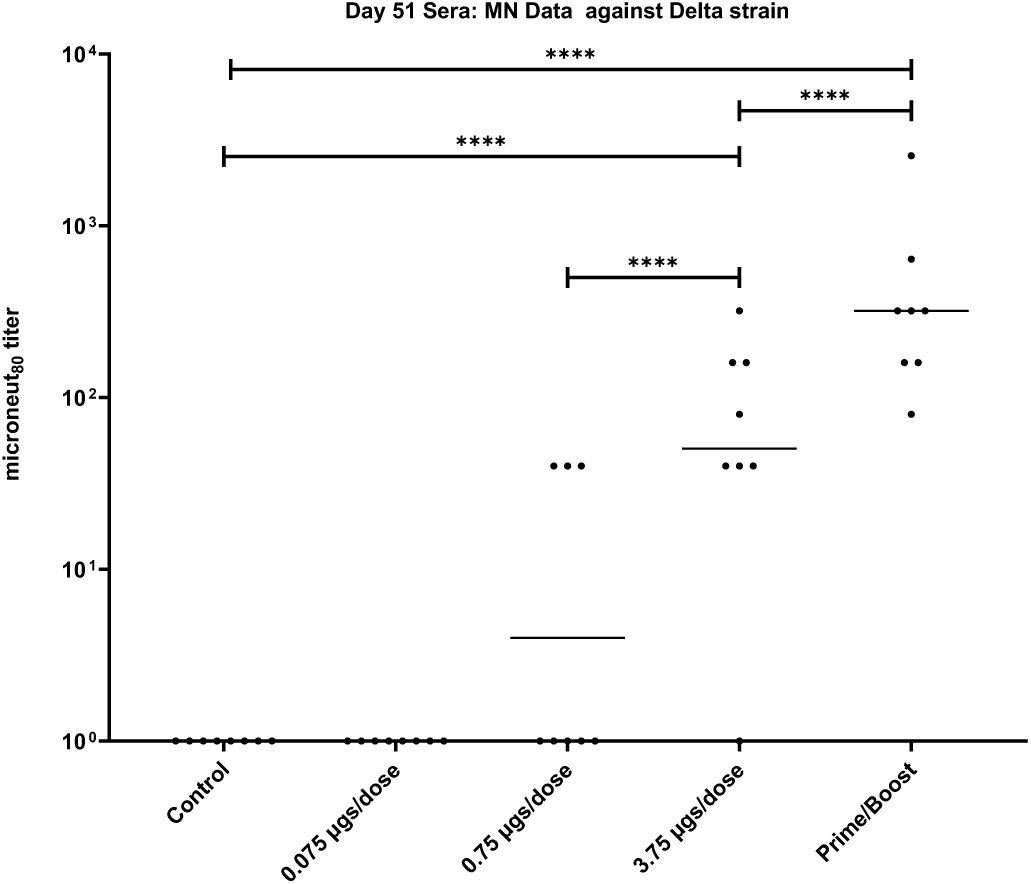
Microneut_80_ data for Day 51 sera from nonhuman primates vaccinated with different doses of SARS-CoV-2 PsIV vaccine. Horizontal bars represent the geometric mean for each group. Statistical significance between groups is denoted by * p ≤ 0.05, ** p ≤ 0.01, *** p ≤ 0.001, **** p ≤ 0.0001.

**Table 1.** Dosage for SARS-CoV-2 PsIV evaluation in nonhuman primates.

| Groups | Vaccine formulation | SARS-CoV-2 PsIV dose (expressed as spike protein equivalent) given |
| --- | --- | --- |
| **Group 1** | Advax-2 + PBS (Controls)<br>(Days 0 and 30) | NA |
| **Group 2** | SARS-CoV-2 PsIV + Advax-2<br>(Days 0 and 30) | 0.075 µg of spike protein/dose |
| **Group 3** | SARS-CoV-2 PsIV + Advax-2<br>(Days 0 and 30) | 0.750 µg of spike protein/dose |
| **Group 4** | SARS-CoV-2 PsIV + Advax-2<br>(Days 0 and 30) | 3.75 µg of spike protein/dose |
| **Group 5** | Plasmid DNA (Days -30 and 0)<br>SARS-CoV-2 PsIV + Advax-2 (Day 30) | 5 mg DNA/dose, and<br>3.75 µg of spike protein/dose |

### Neuroprotective Effects of Vaccination

The neuroprotective potential of vaccination to ameliorate the impact of SARS-CoV-2 infection on brain function was investigated in 20 animals due in part to the fact that the ABSL3 facility accommodates a maximum of 20 NHPs at one time. As such, four animals from each group in Table 1 were challenged with 2 × 10^5^ PFU of SARS-CoV-2 Delta strain on day 111 via intranasal instillation (1 × 10^5^ PFU per nostril).

Approximately 41 days after all monkeys were inoculated with live virus, resting state (RS) MEG scans were acquired in the five groups of subjects. As this is a cross-sectional feasibility study, a single MEG scan was recorded for each subject and a between-subjects design was employed to compare the neurological effects of vaccination following COVID exposure across groups. One of the subjects from the control group was omitted from the study due to an incomplete MRI. SAM statistical parametric maps (SPMs) were constructed for the canonical frequency bands of delta (DC-4 Hz, *δ*), theta (4-8 Hz, θ), alpha (8-13 Hz, *α*), beta (13-30 Hz, *β*), gamma (30-80 Hz, *γ*), and for the full bandwidth (DC-80 Hz, F), for the remaining animals (n=19). Source series (virtual electrodes) were constructed for 42 regions of interest (ROIs; see Methods for list) for each subject based on prior reports of the neurological effects of COVID-19, and power spectral densities were constructed from the source series for each ROI.

Figure 2 depicts the SAM RS SPMs for two representative animals and their associated virtual electrodes while the subjects were lightly anesthetized under propofol to demonstrate the observable effects at an individual level of vaccination for the preservation of neural activity. Panels a and b correspond to an animal from the prime-boost group (group 5) and panels c and d correspond to an unvaccinated control subject (group 1). The SAM SPMs show that the unvaccinated monkey had reduced noise-normalized neuromagnetic power at all bandwidths in comparison to the vaccinated animal after both had been exposed to SARS-CoV-2 B.1.617.2. The source series for the vaccinated subject exhibited widespread synchrony and slow oscillatory activity such as epochs of K-complexes and delta waves, examples of which are visible at ∼70 and 71 s, respectively, in Figure 2b, as well as vertex waves (not shown). In contrast, the virtual electrodes for the unvaccinated subject (Figure 2d) did not appear to be widely synchronized and lacked identifiable waveforms, the absence of which is indicative of encephalopathy.

**Figure 2.**
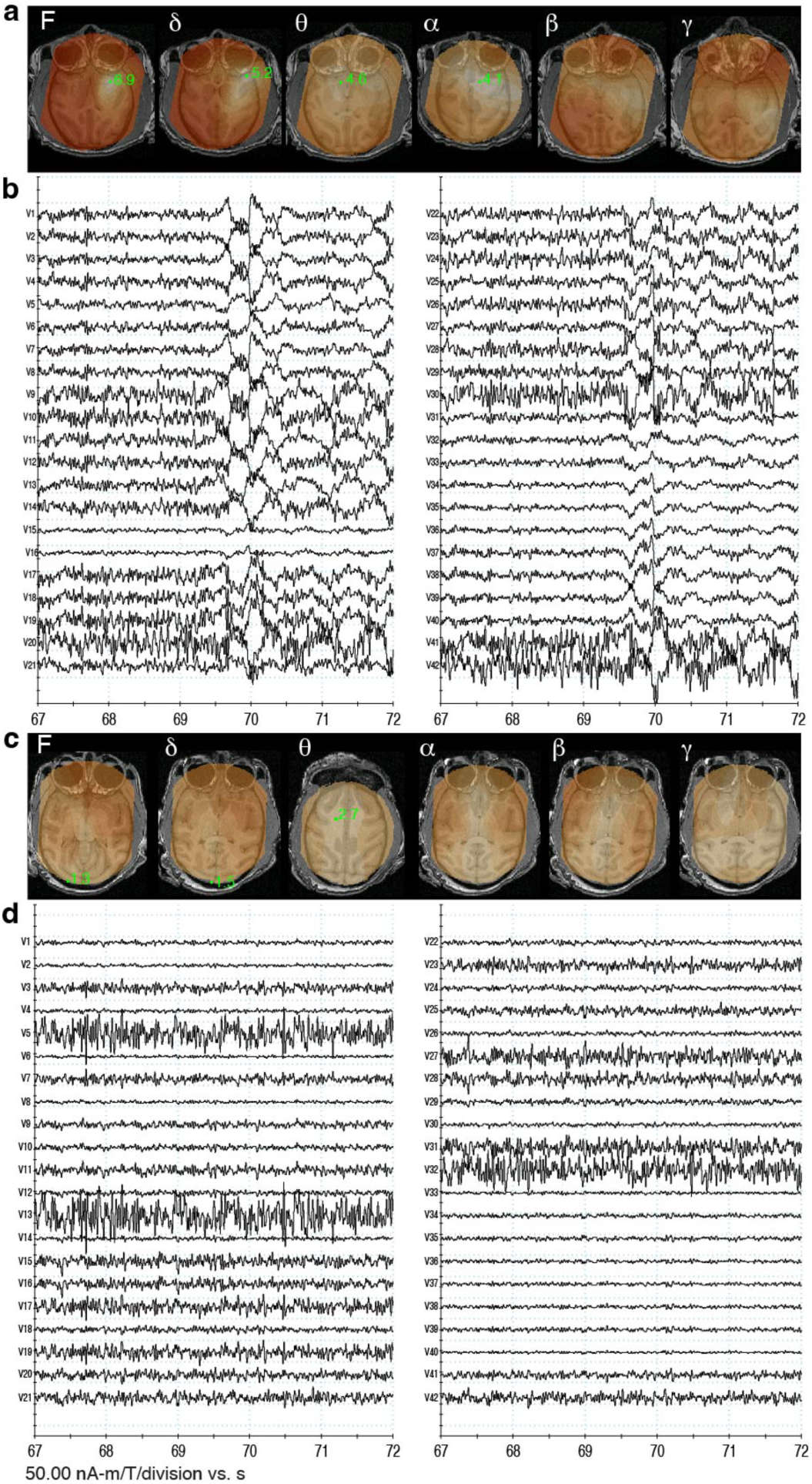
Example SAM resting state maps and MEG virtual electrodes for a vaccinated and unvaccinated subject. **a.** Example axial slices of the SAM SPMs (voxel size = 500 μm^3^) for the full bandwidth as well as for the canonical frequency bands in a representative vaccinated subject. Green dot and number indicate a peak (local maximum) in the SPM plus the associated Ƶ-score. Peaks localize to underlying generators of brain activity. **b.** Virtual electrode traces for the 42 ROIs (shown filtered at 1-40 Hz for viewing purposes) for a 5 s window for the vaccinated subject. **c.** SAM maps for an unvaccinated subject. **d.** Virtual electrodes for the unvaccinated subject. All maps are in radiological coordinates.

Figure 3 depicts example virtual electrode power spectral densities (PSDs) and 95% confidence intervals constructed for four bilateral ROIs from the two representative subjects. Relative to the vaccinated subject (blue), several ROIs for the unvaccinated subject exhibited decreased power across the entire bandwidth.

**Figure 3.**
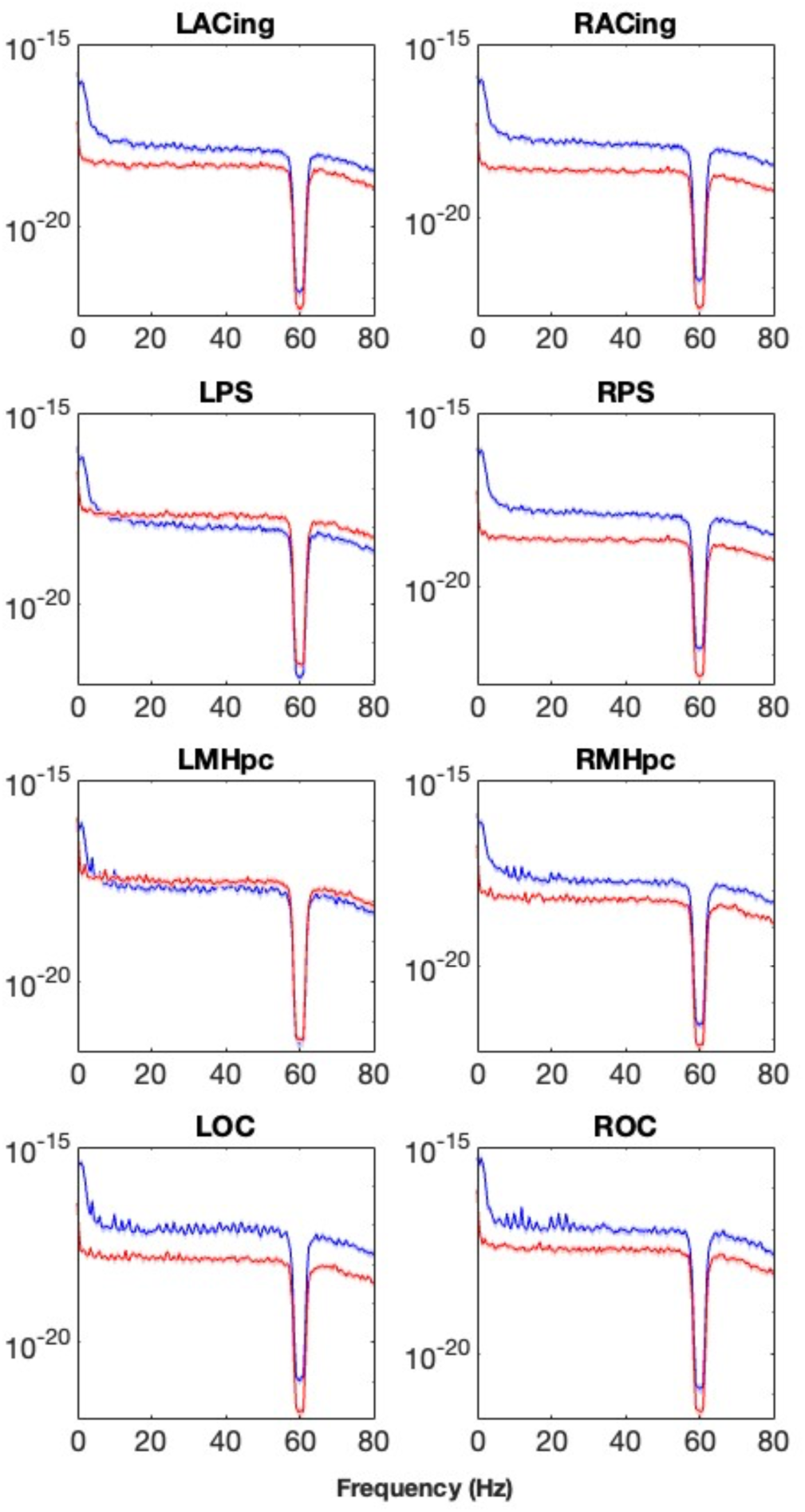
Four bilateral pairs of power spectral densities and 95% confidence intervals for select ROIs for a vaccinated (blue) and unvaccinated (red) subject. PSDs for left and right anterior cingulate (ACing), principal sulcus (PS), middle hippocampus (MHpc), and olfactory cortex (OC) virtual electrodes were calculated for a bandwidth of DC-80 Hz; MEG data were powerline filtered prior to virtual electrode construction. The y-axes are in units of T^2^/Hz.

Figure 4 depicts the average virtual electrode power + standard error of the mean (SEM) per frequency band for the control group and for each of the vaccine groups for the same representative brain areas depicted in figure 3 and Supplementary Figure 1 depicts the average spectral power + SEM per frequency band pooled across all ROIs for each of the groups to illustrate the effects of vaccination on whole brain activity. In particular, the prime-boost group exhibited consistently greater spectral power across all frequency bands for nearly all ROIs. Indeed, multiple linear regression models constructed for the spectral power at each frequency band indicated significant omnibus effects for vaccine group, ROI, and the interaction between vaccine group and ROI, with all p’s < 2.2 x 10^-16^. Given the significant main effects of vaccine group on spectral power, Tukey HSD post hoc tests were conducted within each bandwidth to determine which of the vaccine groups were significantly different from each other for a particular frequency band, and with an overall family-wise error rate of 0.05. The results for these post hoc tests are presented in Table 2 and indicated that the prime boost group had consistently greater spectral power than the controls and the other vaccine groups at all frequency bands.

**Figure 4.**
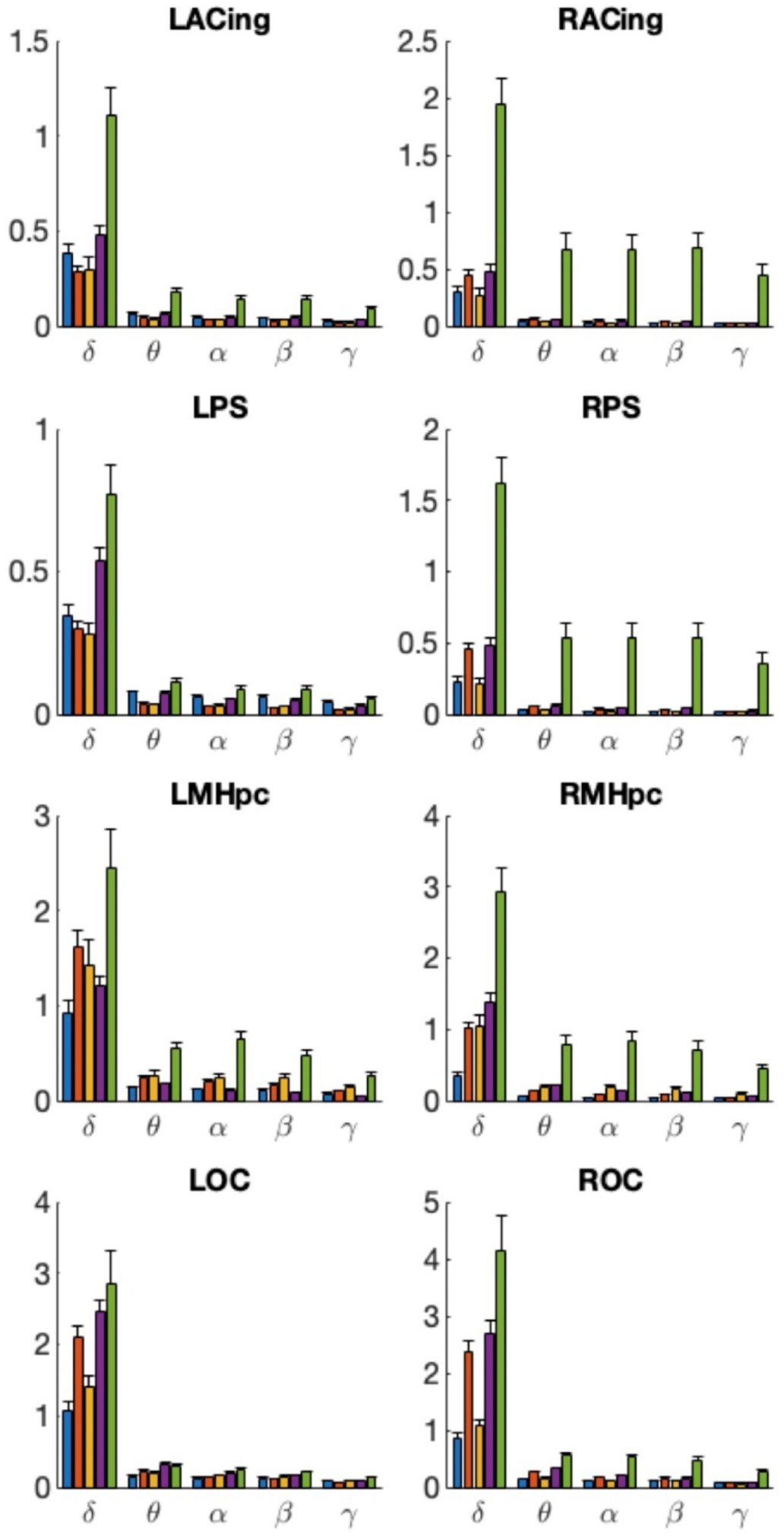
Average spectral power + SEM per frequency band for example ROIs for control and vaccine groups. Blue bars, group 1; group 2, orange; group 3, yellow; group 4, purple; group 5, green. Same ROIs as in Figure 3. The y-axes are in units of 10^-17^ T^2^.

**Table 2.** Tukey HSD contrasts between vaccine groups for each bandwidth. Group 1, controls; 2, 0.075 µg PsIV; 3, 0.75 µg PsIV; 4, 3.75 µg PsIV; 5, prime boost. Entries are p-values, a ‘-‘ denotes insignificance.

| Groups | $\delta$ | $\theta$ | $\alpha$ | $\beta$ | $\gamma$ |
| --- | --- | --- | --- | --- | --- |
| **1-2** | 2E-16 | - | - | - | 0.0026 |
| **1-3** | 1E-05 | - | - | - | - |
| **1-4** | 2E-16 | - | - | - | 0.0053 |
| **1-5** | 2E-16 | 2E-16 | 0 | 2E-16 | 2E-16 |
| **2-3** | 0.0111 | - | 0.0279 | 0.0044 | 0.0202 |
| **2-4** | 0.0154 | - | - | - | - |
| **2-5** | 2E-16 | 2E-16 | 2E-16 | 2E-16 | 2E-16 |
| **3-4** | 1.E-09 | - | - | 0.0195 | 0.0378 |
| **3-5** | 2E-16 | 2E-16 | 2E-16 | 2E-16 | 2E-16 |
| **4-5** | 2E-16 | 2E-16 | 2E-16 | 2E-16 | 2E-16 |

Given the significant interaction term, simple effects contrasts were performed with a Holm correction for the FWER such that the spectral power per bandwidth was contrasted for each of the vaccine groups at each ROI. Table 3 depicts examples of these contrasts for the four representative bilateral brain regions, and Supplementary Table 1 depicts the full set of contrasts for all 42 ROIs. Significant differences (p < 0.05 to p < 2.2 x 10^-16^) in resting state band power were detected in all of the ROIs for the different vaccine groups following infection with SARS-CoV-2 Delta variant. The greatest differences were detected when each of the vaccine groups were compared to either group 1 (unvaccinated) or group 5 (prime boost). Specifically, significantly lower RS activity was observed across the brain at all bandwidths in the unvaccinated group relative to the prime boost group 41 days post-infection. In general, the prime boost group exhibited significantly higher power across the brain relative to the other vaccine groups. It also appears that as the PsIV concentration was increased, the observed power was higher, with the prime boost group having the greatest power.

**Table 3.**
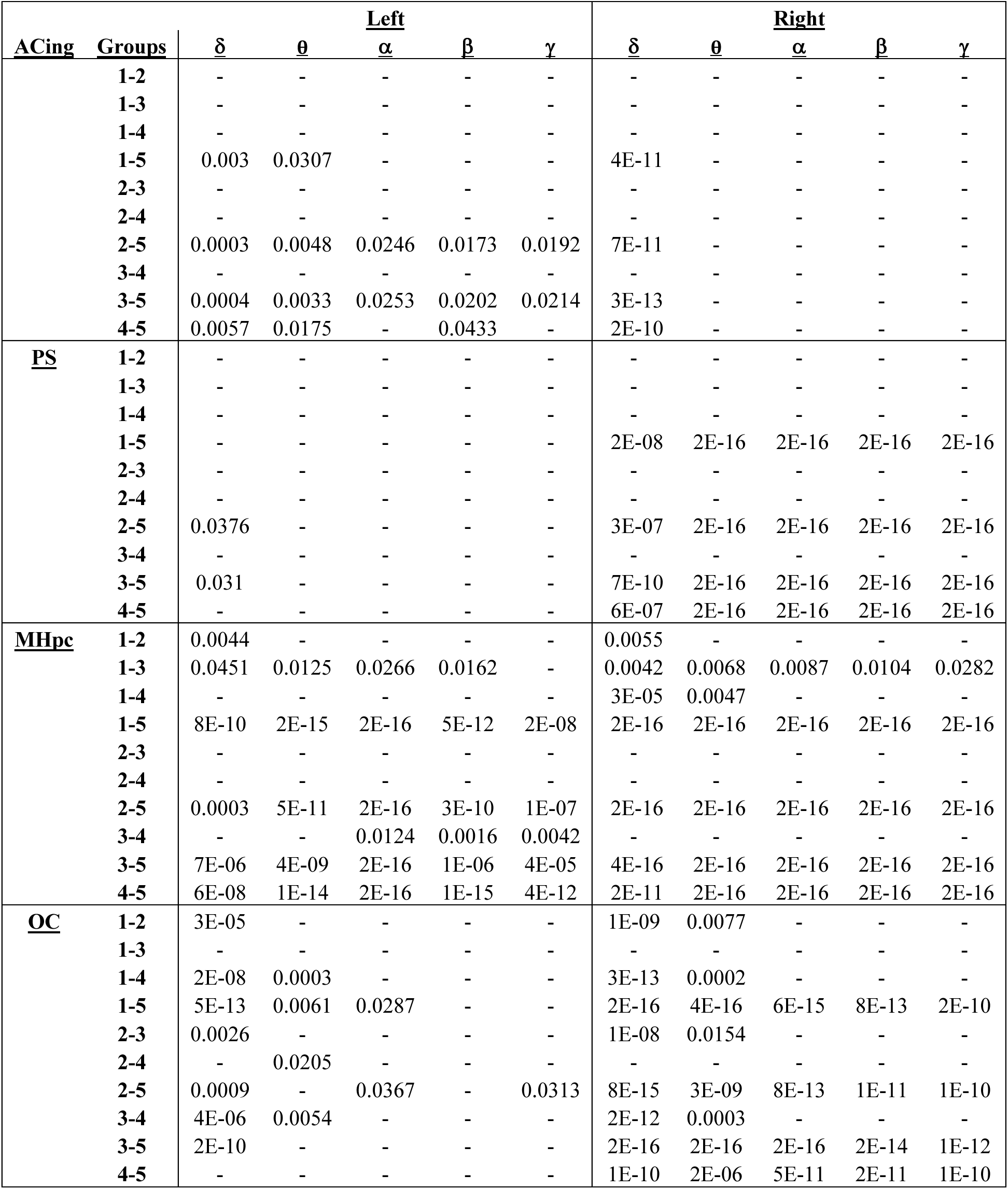
Simple effects contrasts between vaccine groups at the level of example ROIs. Group 1, controls; group 2, 0.075 µg PsIV; group 3, 0.75 µg PsIV; group 4, 3.75 µg PsIV; group 5, prime boost. Entries are p- values, a ‘-‘ denotes insignificance. ACing, anterior cingulate; PS, principal sulcus; MHpc, middle hippocampus; OC, olfactory cortex.

To assess dose-dependent effects of increasing PsIV concentration on total spectral power, a second multiple linear regression model was constructed in which the total power across the full bandwidth was modeled as a numeric, and not categorical, function of the PsIV concentration, ROI, and the interaction between ROI and PsIV concentration. (The prime boost group was omitted from this analysis because it is a combination of two vaccines.) There were significant omnibus effects for vaccine concentration (p < 0.001), ROI (p < 2.2 x 10^-16^), and the interaction between vaccine concentration and ROI (p < 2.2 x 10^-16^). Importantly, the estimated regression coefficient for the PsIV concentration was 0.81, indicating that for every 1 μg increase in PsIV concentration, the full bandwidth spectral power increased 0.81 units. However, in the original regression model of full bandwidth spectral power in which the prime boost group was included as a categorical variable, the estimated regression coefficient for the prime boost vaccine was 18.38, indicating an exceptionally strong effect on spectral power. As a caveat, it remains unclear if some of the differences in brain activity between the vaccine groups were present at baseline because the data were acquired after infection.

## DISCUSSION

COVID-19 is associated with neurobiological deficits in a large percentage of patients, many of whom exhibit abnormal neuroimaging results after a positive diagnosis including abnormal background EEG findings in 96% of patients who receive continuous EEG^17,18,20,27,28^. This study demonstrates the feasibility of MEG as a sensitive tool for detecting changes in brain function. MEG scans demonstrated substantial preservation of neural activity across multiple brain regions in vaccinated subjects compared to unvaccinated controls following viral challenge. The results from our cross-sectional study are consistent with reports applying EEG to measure brain activity in human patients diagnosed with COVID-19^29^, but unlike EEG, MEG possesses sub-mm^3^ spatial resolution, enabling the localization of brain activity from specific cortical and subcortical brain regions^23,30–32^. Animal models, particularly nonhuman primates, are an important component of pre-clinical efforts to understand pathogenesis of SARS-CoV-2 and are critical for studying viral/host factors, disease transmission and pathogenicity, and for testing promising antiviral drugs and vaccine candidates. Macaque monkeys have a long history as subjects in human infectious disease studies^33^, most recently to investigate the neurobiological consequences of COVID-19 infection^34^, and findings in the macaque model are very likely to correspond to findings in humans. An advantage of the nonhuman primate (NHP) model is the ability to track the neurobiological consequences of COVID infection from a known exposure date in a controlled environment in treatment-naïve subjects. Using this model, we identified changes early after vaccination and challenge with live virus. Our group has previously applied MEG to measure RS and sensory gating (SG) brain activity in multiple NHP species and treatment conditions, mapping the impact of alcohol consumption and optogenetic brain stimulation on cortical and subcortical brain regions that co-occur with neuronal activity in vervet and rhesus monkeys^23,35,36^.

As seen in figure 2, the example prime boost vaccinated subject exhibited characteristic resting state brain rhythms under propofol, including widely synchronized activity, vertex waves, K-complexes, and delta waves, all of which are consistent with light/moderate sedation^37^ and all of which occur in humans. In contrast, the unvaccinated subject lacked the observable and normal electrographic architecture under propofol sedation.

While individual susceptibility to anesthesia varies widely^38^, this animal was unlikely to have been more deeply sedated than the vaccinated subject since all subjects in this study were maintained at the same μg/kg/min concentration of propofol and the source series of the control subject did not show signs of slowing or burst suppression. Likewise, the lack of synchrony in the unvaccinated subject’s case is unlikely due to being less sedated than the vaccinated subject because the PSD slopes do not appear to differ between the two subjects. If the unvaccinated subjects were less sedated, low frequency power should decrease and higher frequency power should increase, changing the PSD slopes^39^. Instead, the PSD curves are simply shifted downward, indicating less total power across the spectrum. In summary, the lack of normal propofol sedation architecture in the source series and a full bandwidth decrease in power for the unvaccinated subject in comparison to the prime boost vaccinated subject indicates a disruption in brain function consistent with encephalopathy, likely due to COVID-19 infection.

As a group, the unvaccinated control subjects exhibited the lowest spectral power, likely as a result of COVID- 19-induced encephalopathy. As the PsIV concentration increased, full bandwidth spectral power increased across the ROIs, suggesting that PsIV vaccination conferred dose-dependent neural protection of resting state brain rhythms. The greatest effect was evident for the heterologous prime boost vaccination group which exhibited the greatest spectral power across ROIs in comparison to all other vaccination groups.

Post-COVID Syndrome or Long-COVID occurs in individuals with a history of SARS-CoV-2 infection in which several symptoms last more than three months after infection, including brain fog or cognitive deficits. The COVID and Cognition (COVCOG) study reported that cognitive deficits are one of the most commonly reported symptoms in Long COVID^40^. Recent evidence suggests that changes in resting state brain function in part underlie these deficits and COVID-19 patients exhibit altered functional connectivity that is associated with disease severity^41,42^. Our results extend those findings by demonstrating COVID-19-related reductions in resting state brain activity and further, that vaccination protects against those deficits.

The strong relationship between neurologic protection and vaccination in an NHP model does not specifically address the pathophysiology resulting in potential encephalopathic changes. The host immune response to SARS-CoV-2 infection, including the “cytokine storm” may be one mechanism by which COVID-19 results in neurologic and cognitive sequelae. Such T helper (TH)-1 cytokines as IL-1*β*, IL-6, TNF-*α* and TH-2 cytokines, including IL-4, IL-10 are elevated in serum of COVID-19 patients^43^.

While evidence for direct CNS invasion of SARS-CoV-2 virus as a primary cause of neurologic symptoms remains equivocal, there is some suggestion that symptoms of COVID-19 may be attributable to neurotropic mechanisms and these findings would support that SARS-CoV-2 does invade the CNS. Studies have detected low viral loads in brain tissue early in the infection process though little evidence of inflammation or direct viral cytopathology was detected^13^. Instances of direct CNS invasion have been noted, but the incidence is rare^44^. In contrast, brain lysates from COVID-19 patients who succumbed to the disease demonstrated activation of TGF-*β* signaling, and pathways causing tau hyperphosphorylation typically associated with Alzheimer’s disease (AD)^45^. Diffuse neural inflammatory markers were found in >80% of brains of COVID-19 patients, including microglia in brainstem and hippocampus^46^. Evidence is emerging that proteins and/or RNA of SARS-CoV-2 enter the brain^34,46–48^, and SARS-CoV-2 RNA copies have been detected in olfactory tubercle, medulla, and cerebellum and quantified by RT-qPCR^47^. A recent report suggests the possibility that SARS-CoV-2 Spike protein or its fragments may be released during infection and migrate to CNS and other tissues, resulting in cognitive dysfunction. Infusion of Spike protein into the brain of mice impacts cognitive function^49^. Additionally, RNA from two HCoV strains (229E, OC43) was detected in human brain samples collected at autopsy. Evidence for neurotropism, or affinity for neural tissue, has been documented for coronaviruses such as SARS-CoV and MERS-CoV^50^. Indeed, evidence suggests that SARS-CoV-2 spike proteins may alter permeability of the blood brain barrier by activating the proinflammatory response of brain endothelial cells, thus providing an avenue for entry into the CNS^51,52^. A recent study in nonhuman primates demonstrated immunoreactivity to SARS-CoV-2 nucleocapsid (N) protein in the frontal lobe of rhesus macaques inoculated with *SARS-CoV-2 2019-nCoV/USAWA1/2020,* providing evidence of the presence of viral proteins in the brain within 7 days^34^. In addition to SARS-CoV-2 N, immunolabeling for spike (Spk) markers was found in multiple areas of the primary olfactory cortex, including the olfactory tubercle, the piriform cortex, and the olfactory pole of the entorhinal cortex suggestive of axonal spread from nasal olfactory epithelium.

One limitation of the current study was the small group size for each vaccine dose. Group size was based in part on capacity of the ABSL-3 facility to accommodate 20 animals. Power calculations demonstrate that the study was adequately powered with 4 animals in each group (see Methods). This was borne out by the results demonstrating the robust neuroprotective potential of vaccination to ameliorate COVID-19-related encephalopathic reductions in resting state brain power. It remains unclear whether the clinically used COVID- 19 vaccines offer neuroprotection because they have not been investigated in this capacity.

Another limitation was the cross-sectional comparison of groups after animals had undergone vaccination/no vaccination and challenge with live virus. The cross-sectional nature of the study was based on the parent project that was designed to determine the efficacy of vaccination to protect against SARS-CoV-2 infection. Hence, the ability to define baseline neural activity for each subject was not possible, although the source series the subjects could be visually inspected for the presence, alteration, or absence of expected resting state rhythms, and it is the alteration in these patterns that drive the group-wise differences in the power spectral densities. Preliminary results confirm that vaccination protects against infection-related encephalopathy and the associated reductions in resting state spectral power. Collectively, these results not only underscore the role of vaccination in mitigating severe neurological outcomes but also highlights the capability of MEG to detect subtle yet significant changes in brain function, even in nonhuman primates, that may be overlooked by other imaging modalities. These findings advance our understanding of vaccine-induced neuroprotection and establish MEG as a powerful tool for monitoring brain function in the context of viral infections.

## METHODS

The experiments reported herein were conducted in compliance with the Animal Welfare Act and in accordance with the principles set forth in *Guide for the Care and Use of Laboratory Animals*^53,54^. The study protocol was reviewed and approved by the Wake Forest University School of Medicine Institutional Animal Care and Use Committee (IACUC) and the U.S. Navy Bureau of Medicine and Surgery (BUMED) in compliance with all applicable federal regulations governing the protection of animals and research. A total of 20 female cynomolgus macaques (Macaca Fascicularis, 4-6 years old and average weight 5.8 kg beginning of the study) were subjects in this study. The animals were housed in indoor pens in groups of 3-4 per pen until they were transferred to the ABSL-3 facility for challenge with live virus at which time they were pair-housed in standard quad caging (0.75 x 1.75 x 1.80 m; Allentown Caging, Allentown, PA) with removable wire-mesh partitions that can be used to separate animals when necessary.

### Preparation and Inactivation of SARS-CoV-2

SARS-CoV-2 PsIV was prepared as published previously^55^. Briefly, SARS-CoV-2 strain nCoV/USA- WA1/2020 was propagated in Vero E6 cell cultures and harvested by centrifugation at 3000 x g for 15 minutes. 2000 mL of the culture supernatant containing SARS-CoV-2 was then treated with benzonase (an enzyme degrading free nucleic acids) to remove host cell nucleic acids in the culture supernatant and the volume reduced to 100 mL (concentrating) and buffer exchanged to 10 mM tris buffer containing 150 mM sodium chloride using 100 K MWCO membrane filter cassettes. Concentrated SARS-CoV-2 virus preparation was mixed with AMT (30 µg of AMT per 1 mL of virus) and the resulting mixture was then treated with long wavelength UV light (λ = 365 nm) for 5 minutes (total energy applied = 1.4454 joules). Complete inactivation of psoralen/UVA-treated SARS-CoV-2 virus was confirmed by its inability to grow in permissive cells (Vero E6 cells) by a two-passage virus amplification test. Briefly, 50 µL aliquots of the inactivated virus were used to infect cultured cells in duplicate. After incubation at 37°C for 5-8 days, cells and culture supernatants were examined for the presence of SARS-CoV-2 antigens by indirect immunofluorescence assay and western blot analysis, respectively. The supernatant from this first culture was then incubated with fresh Vero E6 cells for a second round of amplification and testing. Negative results (indicating the absence of virus specific antigens) in both passages confirmed the complete inactivation of SARS-CoV-2.

### Purification and Characterization of SARS-CoV-2 PsIV

SARS-CoV-2 PsIV was purified by chromatographic methods using Cellufine MAX DexS-VirS resin. Briefly, psoralen-inactivated SARS-CoV-2 in 10 mM tris buffer containing 150 mM sodium chloride was passed through a 25 mL Max DexS-VirS column at a flow rate of 0.5 mL per minute, followed by washing with two column volumes of 10 mM tris buffer containing 150 mM sodium chloride. SARS-CoV-2 PsIV bound to the column resin was then eluted using 10 mM tris-HCl buffer containing 500 mM sodium chloride. Fractions containing SARS-CoV-2 PsIV in this elution buffer, identified by Western blot analysis using an anti-SARS-CoV-2 spike protein antibody, were then pooled together as purified SARS-CoV-2 PsIV, then passed through a buffer exchange column with PBS A stabilizer was then added to the pure SARS-CoV-2 PsIV, filtered through a 0.22 μm filter and stored at −80°C. The stabilizer was comprised of a final concentration of 0.5% recombinant human serum albumin, 2% pluronic F-127 and 15% trehalose. The presence of SARS-CoV-2 antigens in the purified inactivated virus preparation was confirmed by western blot using SARS-CoV-2 specific anti-spike protein, anti-nucleoprotein, and anti-envelope protein antibodies. The resulting product was the purified psoralen-inactivated SARS-CoV-2 vaccine (SARS-CoV-2 PsIV). Purity of SARS-CoV-2 PsIV was assessed by gel electrophoresis followed by silver staining. SARS-CoV-2 PsIV titer was determined by spike protein quantitative ELISA using SARS-CoV-2 (2019-nCoV) Spike ELISA Kit (cat# KIT40591, Sino Biological US Inc., PA) and vaccine doses were prepared based on Spike protein concentration.

### Immunogenicity Assessment of SARS-CoV-2 PsIV Vaccine in Nonhuman Primates

The SARS-CoV-2 PsIV vaccine alone and in a prime-boost regimen using two doses of DNA vaccine followed by boosting with SARS-CoV-2 PsIV were evaluated for immunogenicity in adult, female cynomolgus monkeys as shown in Table 1. Five groups of eight animals each were immunized by intramuscular injection (IM) with two doses of different amounts of SARS-CoV-2 PsIV with 10 mg of Advax-2 adjuvant per dose on days 0 and 30 (not shown). Animals in group 1 (Control group) received Advax-2 adjuvant in PBS on days 0 and 30.

Animals in group 2-4 received different amounts of SARS-CoV-2 PsIV (0.075-3.75 µg of spike protein equivalent per dose) with Advax-2 adjuvant on days 0 and 30. Animals in group 5 (prime-boost group), received DNA vaccine encoding the full length SARS-CoV-2 spike glycoprotein^26^ on days -30 and 0, followed by a boosting dose of SARS-CoV-2 PsIV (3.75 µg spike protein equivalent) on day 30. All animals were bled on days -30, 0, 30, 37, 51, 90, and 120, and serum preparations tested for the presence of anti-SARS-CoV-2 neutralizing antibodies using the microneutralization assays.

The ABSL3 environment in which the live virus challenge procedure occurred accommodates caging for 20 monkeys at one time. On day 103 four animals from each of the groups 2-5 and four animals from the control group (total of 20 animals) were moved to ABSL-3 facility to acclimate in preparation for challenge with live SARS-CoV-2 virus and allowed 7 days to acclimate prior to the live virus challenge. On day 111, these twenty animals were challenged with 2×10^5^ PFU of SARS-CoV-2 Delta strain in 0.5 mL solution, via intranasal instillation (1×10^5^ PFU per nostril). These groups served as subjects in the current MEG study. Blood was drawn from all the challenged animals on days 111 and 126. Nasal swabs and throat swabs were collected from the challenged animals on alternate days starting from day 111 until day 125. Results of these assays are described in^26^.

### SARS-CoV-2 Microneutralization Assay

Anti-SARS-CoV-2 neutralizing antibodies in serum were assayed using a microneutralization test against the SARS-CoV-2 Washington strain and the SARS-CoV-2 Delta variant as published previously^55^. Briefly, 200 TCID50 of SARS-CoV-2 test strain was incubated with two-fold dilutions of serum samples in 96 well plates for 1 hour at 37°C. Vero81 cells (2 x 10^4^) were then added to each well and incubated at 37°C for 84 hours. After 84 hours, the cells were fixed, and SARS-CoV-2 titer was measured by quantitating spike protein using SARS-CoV-2 specific anti-S antibody in a standard ELISA format. The highest serum dilution that resulted in ≥80% reduction in absorbance when compared to control samples (without neutralizing antibodies) was determined as the 80% microneutralization titer (MN80).

### Structural MRI

Prior to the start of the MEG study, all animals were tattooed at the nasion and preauricular positions for placement of MEG fiducials and MRI compatible biolipid rings for subsequent T1 weighted structural MRIs. The tattoos are necessary to position the biolipid rings in the same location as the MEG electrodes in order to subsequently co-register the MEG data with the structural MRI. After shaving and aseptically preparing the sites, a Spaulding Tattoo gun was used and a sterile tattoo needle was used for each monkey. Prior to and following tattoo application, triple antibiotic ointment was applied to the site. T1-weighted 3D MPRAGE MRIs of each subject were acquired on a 3T Siemens Skyra using a 32-channel head coil (Siemens AG, Erlangen, Germany). The MRIs were acquired after the vaccination and viral challenge occurred, at approximately 95 days post-infection.

### Resting state MEG Recording and Analysis

All animals were fasted overnight from food but allowed access to water. On the MEG recording day, animals were sedated with ketamine and transported to the MEG suite via an IACUC and institutionally approved transport route. The animals were removed from the transport box and placed on a cart which was covered with blankets. An angiocatheter was placed into the saphenous vein of one leg for propofol administration. Each monkey received a bolus of propofol (2.0-4.0 mg/kg, i.v.) for intubation. Once the animals were intubated, they were maintained on propofol (∼200μg/kg/min) via syringe pump delivery for the duration of the recording procedure. The animals were covered with warming gel packs to maintain body temperature. Head localization was achieved by placing nasion and pre-auricular fiducial coils over the tattoo locations. Head motion was reduced to negligible levels (< 0.2 mm) under anesthesia.

MEG recordings were obtained using a whole head CTF Systems instrument equipped with 275 first-order axial gradiometer coils and 29 reference sensors housed within a magnetically shielded room (MSR; Vacuumschmelze GmbH & Co.; Hanau, Germany). Five minutes of resting state data were acquired while the subjects were maintained in the supine position. Neuromagnetic responses were sampled at 2400 Hz with a bandwidth of DC-600 Hz. All preprocessing and beamforming were performed in the CTF MEG™ Software package (CTF MEG Neuro Innovations, Inc., Coquitlam, BC, Canada). Data were preprocessed using synthetic 3rd order gradient balancing, whole trial DC offsetting, and band pass filtering from DC-80 Hz with powerline filtering ^23,56,57^. MEG data were then co-registered with the monkey’s anatomical MRI data based on the fiducials. From this, a multiple-overlapping-spheres model of the head and whole brain volume was generated^58^. Whole-brain, resting state, noise-normalized, Ƶ-score synthetic aperture magnetometry (SAM)^31,59^ statistical parametric maps of biomagnetic activity were constructed from the MEG data on a per subject basis at a bandwidth of DC-80 Hz and with a voxel size of 2 mm to ensure coregistration with the MRI. For display purposes (see Figure 2), two representative NHPs (one control and one prime boost) were also beamformed with SAM at delta (DC-4 Hz), theta (4-8 Hz), alpha (8-13 Hz), beta (13-30 Hz), gamma (30-80 Hz), and the full bandwidth (DC-80 Hz), and with a voxel size of 500 μm^3^.

For each animal 42 non-adjacent bilateral regions of interest (ROIs) were manually identified in native brain space on each animal’s MRI by a primate neuroanatomist (J.D.). ROIs were chosen based on existing literature describing the impact of COVID-19 infection on brain volumetric measures^14^, fluorodeoxyglucose uptake, or those areas underlying executive and cognitive function. The list of mainly bilateral ROIs includes: L/R ACing, left/right anterior cingulate cortex; L/R MO, medial orbitofrontal cortex; L/R LO, lateral orbitofrontal cortex; L/R PS, principle sulcus; L/R NAc, nucleus accumbens; L/R Cau, caudate n.; L/R PostPut, posterior putamen; L/R precuneus; L/R Am(lat), lateral amygdala; L/R Am(BM-BL), basomedial/basolateral amygdala; L/R MHpc, middle hippocampus; L/R AHpc, anterior hippocampus; L/R CHpc, central hippocampus; L/R CbA, anterior cerebellum; L/R CbP, posterior cerebellum; midline Vermis; L/R AI, anterior insula; OB, midline olfactory bulb; L/R OB, olfactory bulb; L/R Obpost, posterior olfactory bulb; L/R Obmed, medial olfactory bulb; and L/R OC, olfactory cortex. Virtual electrodes (source series, equivalent to local field potentials, DC-80 Hz bandwidth) were extracted for each ROI, divided into 20 second epochs (15 epochs per animal), and the power spectral densities (PSDs) for each epoch were computed by Welch’s method in Matlab 2024a (Natick, Massachusetts: The MathWorks Inc.). 95% confidence intervals (CIs) were calculated for the PSDs for figure 3 using the mean and standard deviations of the PSDs and critical value of the t-distribution for the appropriate sample size. For all subjects, the PSDs were split into the canonical frequency bands of delta (DC-4 Hz), theta (4-8 Hz), alpha (8-13 Hz), beta (13-30 Hz), and gamma (30-80 Hz), as well as the full bandwidth (DC-80 Hz) and total power was measured within each band.

### Statistics

For the microneutralization results, data analysis was performed using GraphPad Prism 9.0.2 or 9.4.1. Student’s *t*-tests were used to compare the microneutralization data between groups.

For the MEG data, multiple linear regression models were constructed in R version 4.2.0, “Vigorous Calisthenics”^60^, with separate models for spectral power within each of the frequency bands. Each modeled the spectral power for a given bandwidth as a function of vaccine group (groups 1-5), bandwidth (delta, theta, alpha, beta, gamma, full bandwidth), and the interaction of vaccine group and bandwidth. All factors were modeled as indicators, and the power for each of the bandwidths was modeled as a continuous variable. Given significant main effects of vaccine group on spectral power, Tukey HSD post hoc tests were conducted within each bandwidth to determine which of the vaccine groups were significantly different from each other for a particular frequency band, and with an overall family-wise error rate (FWER) of 0.05. Given the significant interaction between vaccine group and brain region, simple effects contrasts were conducted within each bandwidth to determine if power differed between the vaccine groups for each brain ROI. Simple effects contrasts were conducted with the phia package version 0.2-1 in R^61^. The Holm method was used to control the FWER at an overall alpha of 0.05 for the simple effects test; this method is equivalent to the Bonferroni method in controlling Type I error but is better at controlling the Type II error rate^62^.

To assess dose-dependent effects of increasing PsIV concentration on total spectral power, a final multiple linear regression model was constructed in which the total power across the full bandwidth was modeled as a numeric, and not categorical, function of the PsIV concentration, ROI, and the interaction between ROI and PsIV concentration. The ROI factor was modeled as an indicator variable, and the full bandwidth power was modeled as a continuous variable. The prime boost group was omitted from this analysis because it is a combination of two vaccines. Given significant main effects of PsIV vaccine concentration on spectral power, the estimated coefficient for PsIV concentration was extracted to assess the magnitude of dose-dependent vaccine effects on preserving brain function.

Group size was determined using spectral power calculated from the 42 ROIs. An ANOVA was conducted in SPSS including main effects of group (group 1, unvaccinated, vs. group 5, prime boost) as well as time with power in the alpha band as the dependent variable. Groups were significantly different after false discovery rate correction at all but three ROIs. Power analysis was conducted using means and standard deviations from this preliminary analysis for the anterior cingulate (vaccinated mean=1.55, SD=1.74; unvaccinated mean=0.26, SD=0.26) and orbitofrontal cortex (vaccinated mean=6.76, SD=10.68; unvaccinated mean=0.34, SD=0.16).

Using a range of +/- one standard deviation from the mean, statistical power for the ACC ranged from 0.907 to 0.999 and for the orbitofrontal cortex from 0.904 to 0.999. These results suggest a group size of n = 4 was adequately powered to determine direct contrasts between the groups.

## DATA AVAILABILITY

Data will be made available on reasonable request.

## ACKNOWLEDGEMENTS

JD was supported by NS132017, COVID-19 Research Project pilot, Ignition Funds from the Wake Forest School of Medicine Translational Sciences Institute (JBD). DFE, ZL, AKS and KRP were funded by DHP RDT&E supplemental COVID funding. Development of Advax-CpG adjuvant was supported by funding from National Institute of Allergy and Infectious Diseases of the National Institutes of Health to Vaxine Pty, Ltd., under Contracts HHS-N272201400053C, HHS-N272200800039C and U01-AI061142. Additional details on Advax (VO_0005207) and Advax-CpG55.2 (VO_0005324) adjuvants can be found on the NIAID Vaccine Adjuvant Compendium database at https://vac.niaid.nih.gov. Special thanks to Dr. Robert Kotloski for discussions about neurophysiological rhythms under propofol as well as encephalopathic rhythms.

## DISCLAIMER

The views expressed in this article reflect the research conducted by the authors, and do not necessarily reflect the official policy or position of the Department of the Navy, the Department of Defense, or the U.S. Government. Copyright Statement: The authors are military service members or federal/contracted employees of the U.S. Government. This work was prepared as part of their official duties. Title 17 U.S.C. § 105 provides that ‘Copyright protection under this title is not available for any work of the U.S. Government.’ Title 17 U.S.C.

§ 101 defines a U.S. Government work as a work prepared by military service members or employees of the

U.S. Government as part of that persons’ official duties.

## AUTHOR CONTRIBUTIONS

J.S.K. contributed to experimental design, data collection, data analysis, interpretation and manuscript preparation. J.B.D. contributed to experimental design, data collection, data analysis, interpretation and manuscript preparation. J.A.R. contributed to data collection, data analysis, interpretation and manuscript preparation. J.W.S contributed to experimental design, interpretation, and manuscript preparation. D.W.G. contributed to data collection, interpretation, and manuscript preparation. A.T.D. contributed to data collection, and manuscript preparation. P.M.E. contributed to data collection, and manuscript preparation. M.B. contributed to experimental design, data collection, and manuscript preparation. C.S.G contributed to data collection, and manuscript preparation. D.F.E. contributed to experimental design, data collection, data interpretation and manuscript preparation. Z.L. contributed to data collection and data interpretation. A.K.S. contributed to experimental design, data collection data interpretation and manuscript preparation. K.R.P. contributed to experimental design, data interpretation and manuscript preparation. N.P. provided the adjuvant and contributed to manuscript preparation.

## MATERIALS & CORRESPONDENCE

Correspondence and material requests should be addressed to Dr. James Daunais and Dr. John Sanders.

