## Supplementary Figure 1 for "Magnetoencephalography Reveals Neuroprotective Effects of COVID-19 Vaccination in Non-Human Primates"

### ADDITIONAL INFORMATION

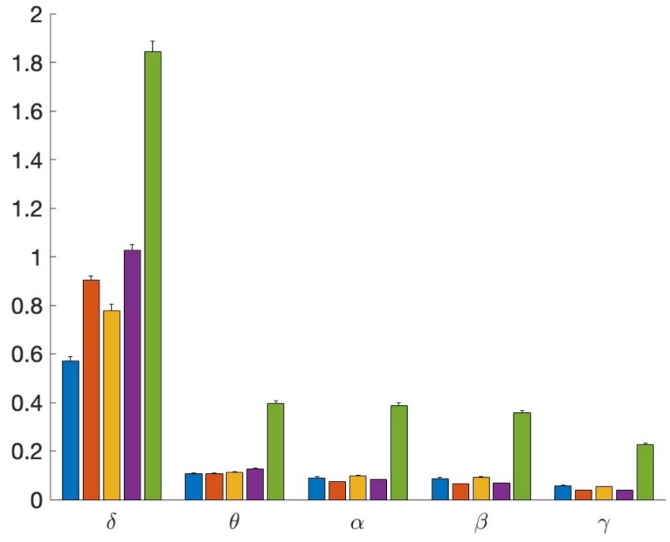


**Supplementary Figure 1.**

### EXTENDED LEGENDS

**Supplementary Figure 1. Average spectral power + SEM per frequency band pooled across all ROIs for control and vaccine groups.** Blue bars, group 1; group 2, orange; group 3, yellow; group 4, purple; group 5, green. The y-axes are in units of $10^{-17}$ $T^2$.
