## Supplementary Table 1 for "Magnetoencephalography Reveals Neuroprotective Effects of COVID-19 Vaccination in Non-Human Primates"

| | | Left | | | | | | Right | | | | | |
|---|---|---|---|---|---|---|---|---|---|---|---|---|---|
| **ACing** | **Groups** | **δ** | **θ** | **α** | **β** | **γ** | **F** | **δ** | **θ** | **α** | **β** | **γ** | **F** |
| | **1-2** | - | - | - | - | - | - | - | - | - | - | - | - |
| | **1-3** | - | - | - | - | - | - | - | - | - | - | - | - |
| | **1-4** | - | - | - | - | - | - | - | - | - | - | - | - |
| | **1-5** | 0.003 | 0.031 | - | - | - | 0.012 | 4E-11 | 2E-16 | 2E-16 | 2E-16 | 2E-16 | 2E-16 |
| | **2-3** | - | - | - | - | - | - | - | - | - | - | - | - |
| | **2-4** | - | - | - | - | - | - | - | - | - | - | - | - |
| | **2-5** | 3E-04 | 0.005 | 0.025 | 0.017 | 0.019 | 0.002 | 7E-11 | 2E-16 | 2E-16 | 2E-16 | 2E-16 | 2E-16 |
| | **3-4** | - | - | - | - | - | - | - | - | - | - | - | - |
| | **3-5** | 4E-04 | 0.003 | 0.025 | 0.02 | 0.021 | 0.002 | 3E-13 | 2E-16 | 2E-16 | 2E-16 | 2E-16 | 2E-16 |
| | **4-5** | 0.006 | 0.018 | - | 0.043 | - | 0.012 | 2E-10 | 2E-16 | 2E-16 | 2E-16 | 2E-16 | 2E-16 |
| **MO** | **1-2** | - | - | - | - | - | - | - | - | - | - | - | - |
| | **1-3** | - | - | - | - | - | - | - | - | - | - | - | - |
| | **1-4** | - | - | - | - | - | - | - | - | - | - | - | - |
| | **1-5** | - | - | - | - | - | - | 6E-11 | 2E-16 | 2E-16 | 2E-16 | 2E-16 | 2E-16 |
| | **2-3** | - | - | - | - | - | - | - | - | - | - | - | - |
| | **2-4** | - | - | - | - | - | - | - | - | - | - | - | - |
| | **2-5** | - | - | - | - | - | 0.048 | 4E-07 | 2E-16 | 2E-16 | 2E-16 | 2E-16 | 2E-16 |
| | **3-4** | 0.025 | - | - | - | - | - | - | - | - | - | - | - |
| | **3-5** | - | 0.034 | - | - | - | 0.04 | 1E-11 | 2E-16 | 2E-16 | 2E-16 | 2E-16 | 2E-16 |
| | **4-5** | - | - | - | - | - | - | 2E-07 | 2E-16 | 2E-16 | 2E-16 | 2E-16 | 2E-16 |
| **LO** | **1-2** | - | 5E-08 | 5E-08 | 6E-09 | 9E-10 | 3E-08 | - | - | - | - | - | - |
| | **1-3** | 0.042 | 7E-08 | 1E-07 | 2E-08 | 3E-09 | 5E-08 | - | - | - | - | - | - |
| | **1-4** | - | 1E-05 | 1E-06 | 2E-07 | 2E-08 | 3E-06 | - | - | - | - | - | - |
| | **1-5** | - | 8E-06 | 6E-06 | 2E-06 | 3E-07 | 1E-05 | 5E-06 | 1E-13 | 4E-15 | 2E-15 | 7E-16 | 5E-16 |
| | **2-3** | - | - | - | - | - | - | - | - | - | - | - | - |
| | **2-4** | - | - | - | - | - | - | - | - | - | - | - | - |
| | **2-5** | - | - | - | - | - | - | 0.001 | 3E-13 | 5E-16 | 2E-16 | 2E-16 | 1E-15 |
| | **3-4** | - | - | - | - | - | - | - | - | - | - | - | - |
| | **3-5** | - | - | - | - | - | - | 9E-07 | 9E-16 | 2E-16 | 2E-16 | 2E-16 | 2E-16 |
| | **4-5** | - | - | - | - | - | - | 0.001 | 1E-12 | 4E-15 | 8E-16 | 2E-16 | 6E-15 |
| **PS** | **1-2** | - | - | - | - | - | - | - | - | - | - | - | - |
| | **1-3** | - | - | - | - | - | - | - | - | - | - | - | - |
| | **1-4** | - | - | - | - | - | - | - | - | - | - | - | - |
| | **1-5** | - | - | - | - | - | - | 2E-08 | 2E-16 | 2E-16 | 2E-16 | 2E-16 | 2E-16 |
| | **2-3** | - | - | - | - | - | - | - | - | - | - | - | - |
| | **2-4** | - | - | - | - | - | - | - | - | - | - | - | - |
| | **2-5** | 0.038 | - | - | - | - | - | 3E-07 | 2E-16 | 2E-16 | 2E-16 | 2E-16 | 2E-16 |
| | **3-4** | - | - | - | - | - | - | - | - | - | - | - | - |
| | **3-5** | 0.031 | - | - | - | - | - | 7E-10 | 2E-16 | 2E-16 | 2E-16 | 2E-16 | 2E-16 |
| | **4-5** | - | - | - | - | - | - | 6E-07 | 2E-16 | 2E-16 | 2E-16 | 2E-16 | 2E-16 |
| **NAc** | **1-2** | 0.005 | - | - | - | - | - | 5E-04 | - | - | - | - | - |
| | **1-3** | 0.015 | - | - | - | - | - | - | - | - | - | - | - |
| | **1-4** | 1E-04 | - | - | - | - | - | 5E-05 | - | - | - | - | - |
| | **1-5** | 3E-09 | 0.04 | 0.017 | 0.009 | 0.009 | 8E-05 | 4E-12 | 3E-15 | 2E-15 | 2E-16 | 2E-16 | 2E-16 |
| | **2-3** | - | - | - | - | - | - | 0.023 | - | - | - | - | - |
| | **2-4** | - | - | - | - | - | - | - | - | - | - | - | - |
| | **2-5** | 9E-04 | 0.029 | 0.026 | 0.012 | 0.011 | 0.002 | 2E-04 | 8E-16 | 4E-15 | 2E-16 | 2E-16 | 2E-16 |
| | **3-4** | - | - | - | - | - | - | 0.003 | - | - | - | - | - |
| | **3-5** | 2E-04 | 0.013 | 0.035 | 0.022 | 0.01 | 0.001 | 2E-09 | 2E-16 | 8E-15 | 2E-16 | 2E-16 | 2E-16 |
| | **4-5** | 0.026 | 0.01 | 0.009 | 0.005 | 0.007 | 0.003 | 0.002 | 1E-15 | 2E-15 | 2E-16 | 2E-16 | 2E-16 |
| **Cau** | **1-2** | - | - | - | - | - | - | - | - | - | - | - | - |
| | **1-3** | - | - | - | - | - | - | - | - | - | - | - | - |
| | **1-4** | - | - | - | - | - | - | - | - | - | - | - | - |
| | **1-5** | 0.006 | 0.016 | 0.009 | 0.008 | 0.012 | 0.003 | 5E-10 | 2E-16 | 2E-16 | 2E-16 | 2E-16 | 2E-16 |

| | | | | | | | | | | | | | |
|---|---|---|---|---|---|---|---|---|---|---|---|---|---|
| | **2-3** | - | - | - | - | - | - | - | - | - | - | - | - |
| | **2-4** | - | - | - | - | - | - | - | - | - | - | - | - |
| | **2-5** | 4E-04 | 8E-06 | 6E-05 | 8E-05 | 1E-04 | 1E-05 | 2E-08 | 2E-16 | 2E-16 | 2E-16 | 2E-16 | 2E-16 |
| | **3-4** | - | - | - | - | - | - | - | - | - | - | - | - |
| | **3-5** | 5E-05 | 3E-06 | 4E-05 | 6E-05 | 5E-05 | 3E-06 | 7E-13 | 2E-16 | 2E-16 | 2E-16 | 2E-16 | 2E-16 |
| | **4-5** | 4E-04 | 5E-06 | 3E-05 | 3E-05 | 5E-05 | 5E-06 | 2E-09 | 2E-16 | 2E-16 | 2E-16 | 2E-16 | 2E-16 |
| **PostPut** | **1-2** | - | 1E-15 | 2E-16 | 2E-16 | 2E-16 | 2E-15 | - | - | - | - | - | - |
| | **1-3** | - | 5E-15 | 5E-16 | 2E-16 | 2E-16 | 9E-15 | - | - | - | - | - | - |
| | **1-4** | - | 1E-14 | 2E-16 | 2E-16 | 2E-16 | 2E-15 | - | - | - | - | - | - |
| | **1-5** | - | 2E-07 | 3E-09 | 4E-10 | 6E-12 | 1E-06 | 4E-12 | 2E-16 | 2E-16 | 2E-16 | 2E-16 | 2E-16 |
| | **2-3** | - | - | - | - | - | - | - | - | - | - | - | - |
| | **2-4** | - | - | - | - | - | - | - | - | - | - | - | - |
| | **2-5** | 3E-04 | 0.003 | 0.006 | 0.007 | 0.01 | 9E-04 | 1E-08 | 2E-16 | 2E-16 | 2E-16 | 2E-16 | 2E-16 |
| | **3-4** | - | - | - | - | - | - | - | - | - | - | - | - |
| | **3-5** | 8E-05 | 0.005 | 0.019 | 0.027 | 0.031 | 0.002 | 2E-11 | 2E-16 | 2E-16 | 2E-16 | 2E-16 | 2E-16 |
| | **4-5** | 5E-04 | 0.008 | 0.008 | 0.008 | 0.009 | 0.001 | 5E-08 | 2E-16 | 2E-16 | 2E-16 | 2E-16 | 2E-16 |
| **Precuneus** | **1-2** | - | - | - | - | - | - | - | - | - | - | - | - |
| | **1-3** | - | - | - | - | - | - | - | - | - | - | - | - |
| | **1-4** | - | - | - | - | - | - | - | - | - | - | - | - |
| | **1-5** | - | - | - | - | - | - | 0.049 | 0.018 | 0.018 | - | - | 0.027 |
| | **2-3** | - | - | - | - | - | - | 0.019 | - | - | - | - | - |
| | **2-4** | - | - | - | - | - | - | - | - | - | - | - | - |
| | **2-5** | 0.027 | 0.004 | 0.004 | 0.014 | 0.016 | 0.006 | 0.005 | 5E-04 | 6E-04 | 0.003 | 0.004 | 0.0007 |
| | **3-4** | - | - | - | - | - | - | - | - | - | - | - | - |
| | **3-5** | - | - | - | - | - | - | - | 0.038 | 0.033 | - | - | - |
| | **4-5** | - | 0.008 | 0.006 | 0.016 | 0.018 | 0.01 | 0.027 | 0.001 | 0.001 | 0.005 | 0.005 | 0.002 |
| **Am(lat)** | **1-2** | 0.009 | - | - | - | - | - | 0.002 | - | - | - | - | - |
| | **1-3** | 0.031 | - | - | - | - | - | - | - | - | - | - | - |
| | **1-4** | 0.018 | - | - | - | - | - | 8E-06 | 0.004 | - | - | - | 0.0262 |
| | **1-5** | 1E-05 | 0.004 | 0.005 | - | - | 0.005 | 4E-13 | 2E-16 | 2E-16 | 2E-16 | 2E-16 | 2E-16 |
| | **2-3** | - | - | - | 0.048 | - | - | 0.007 | - | - | - | - | - |
| | **2-4** | - | - | - | - | - | - | - | - | - | - | - | - |
| | **2-5** | - | 0.003 | 2E-04 | 0.002 | 0.004 | 0.002 | 5E-06 | 2E-16 | 2E-16 | 2E-16 | 2E-16 | 2E-16 |
| | **3-4** | - | - | - | - | - | - | 2E-05 | 0.012 | - | - | - | - |
| | **3-5** | 0.016 | - | - | - | - | - | 5E-13 | 2E-16 | 2E-16 | 2E-16 | 2E-16 | 2E-16 |
| | **4-5** | 0.03 | 0.038 | 5E-04 | 0.003 | 0.002 | 0.001 | 0.003 | 2E-11 | 2E-16 | 2E-16 | 2E-16 | 2E-16 |
| **Am(BM-BL)** | **1-2** | 3E-05 | - | - | - | - | - | 7E-04 | - | - | - | - | - |
| | **1-3** | - | - | - | - | - | - | - | - | - | - | - | - |
| | **1-4** | 4E-04 | 0.031 | - | - | - | - | 1E-11 | 7E-05 | - | - | - | 0.0033 |
| | **1-5** | 1E-06 | 0.003 | 0.004 | - | - | 0.003 | 2E-16 | 2E-16 | 2E-16 | 2E-16 | 2E-16 | 2E-16 |
| | **2-3** | 1E-04 | - | - | - | - | - | 0.001 | - | - | - | - | - |
| | **2-4** | - | - | - | - | - | - | 3E-04 | 0.01 | - | - | - | - |
| | **2-5** | - | 0.043 | 0.001 | 0.009 | 0.009 | 0.014 | 2E-16 | 2E-16 | 2E-16 | 2E-16 | 2E-16 | 2E-16 |
| | **3-4** | 0.001 | - | - | - | - | - | 5E-12 | 3E-04 | - | - | - | 0.0042 |
| | **3-5** | 4E-06 | 0.009 | 0.006 | 0.048 | 0.027 | 0.001 | 2E-16 | 2E-16 | 2E-16 | 2E-16 | 2E-16 | 2E-16 |
| | **4-5** | - | - | 0.005 | 0.011 | 0.004 | 0.008 | 3E-10 | 6E-14 | 2E-16 | 2E-16 | 2E-16 | 2E-16 |
| **MHpc** | **1-2** | 0.004 | - | - | - | - | - | 0.006 | - | - | - | - | - |
| | **1-3** | 0.045 | 0.013 | 0.027 | 0.016 | - | 0.017 | 0.004 | 0.007 | 0.009 | 0.01 | 0.028 | 0.0041 |
| | **1-4** | - | - | - | - | - | - | 3E-05 | 0.005 | - | - | - | 0.0226 |
| | **1-5** | 8E-10 | 2E-15 | 2E-16 | 5E-12 | 2E-08 | 1E-13 | 2E-16 | 2E-16 | 2E-16 | 2E-16 | 2E-16 | 2E-16 |
| | **2-3** | - | - | - | - | - | - | - | - | - | - | - | - |
| | **2-4** | - | - | - | - | - | - | - | - | - | - | - | - |
| | **2-5** | 3E-04 | 5E-11 | 2E-16 | 3E-10 | 1E-07 | 6E-10 | 2E-16 | 2E-16 | 2E-16 | 2E-16 | 2E-16 | 2E-16 |
| | **3-4** | - | - | 0.012 | 0.002 | 0.004 | 0.008 | - | - | - | - | - | - |
| | **3-5** | 7E-06 | 4E-09 | 2E-16 | 1E-06 | 4E-05 | 5E-08 | 4E-16 | 2E-16 | 2E-16 | 2E-16 | 2E-16 | 2E-16 |
| | **4-5** | 6E-08 | 1E-14 | 2E-16 | 1E-15 | 4E-12 | 4E-16 | 2E-11 | 2E-16 | 2E-16 | 2E-16 | 2E-16 | 2E-16 |

| | | | | | | | | | | | | | |
|---|---|---|---|---|---|---|---|---|---|---|---|---|---|
| **AHpc** | **1-2** | 0.007 | - | - | - | - | - | 0.01 | - | - | - | - | - |
| | **1-3** | 0.005 | 9E-04 | 0.004 | 0.003 | 0.017 | 0.002 | 0.049 | - | - | - | - | 0.0405 |
| | **1-4** | - | - | - | - | - | - | 5E-06 | 0.002 | - | - | - | 0.0144 |
| | **1-5** | 1E-09 | 2E-11 | 2E-16 | 1E-08 | 5E-06 | 2E-10 | 2E-16 | 2E-16 | 2E-16 | 2E-16 | 2E-16 | 2E-16 |
| | **2-3** | - | - | 0.029 | 0.007 | 0.017 | 0.037 | - | - | - | - | - | - |
| | **2-4** | - | - | - | - | - | - | 0.029 | 0.036 | - | - | - | - |
| | **2-5** | 3E-04 | 3E-08 | 2E-16 | 2E-08 | 2E-06 | 2E-08 | 3E-12 | 2E-16 | 2E-16 | 2E-16 | 2E-16 | 2E-16 |
| | **3-4** | - | 0.021 | 0.002 | 2E-04 | 7E-04 | 0.001 | 0.005 | - | - | - | - | - |
| | **3-5** | 5E-04 | 3E-04 | 1E-09 | 0.004 | 0.019 | 4E-04 | 2E-14 | 2E-16 | 2E-16 | 2E-16 | 2E-16 | 2E-16 |
| | **4-5** | 1E-06 | 3E-09 | 2E-16 | 3E-11 | 9E-09 | 9E-12 | 1E-06 | 2E-16 | 2E-16 | 2E-16 | 2E-16 | 2E-16 |
| **CHpc** | **1-2** | 0.001 | 0.032 | - | - | - | 0.024 | 0.021 | - | - | - | - | - |
| | **1-3** | 0.024 | 0.016 | 0.026 | 0.019 | 0.049 | 0.013 | 0.002 | 0.003 | 0.003 | 0.007 | 0.017 | 0.0019 |
| | **1-4** | - | - | - | - | - | - | 0.002 | 0.025 | - | - | - | - |
| | **1-5** | 7E-14 | 2E-16 | 2E-16 | 5E-14 | 4E-10 | 2E-16 | 1E-11 | 1E-08 | 1E-08 | 1E-06 | 2E-05 | 2E-09 |
| | **2-3** | - | - | - | - | - | - | - | - | 0.025 | 0.038 | - | - |
| | **2-4** | 0.002 | - | 0.042 | 0.043 | - | 0.01 | - | - | - | - | - | - |
| | **2-5** | 3E-06 | 5E-14 | 2E-16 | 2E-10 | 1E-07 | 6E-11 | 1E-06 | 2E-06 | 2E-07 | 1E-05 | 8E-05 | 5E-07 |
| | **3-4** | 0.034 | 0.049 | 0.016 | 0.005 | 0.01 | 0.005 | - | - | - | - | - | - |
| | **3-5** | 2E-08 | 4E-13 | 2E-16 | 2E-08 | 4E-06 | 3E-10 | 7E-05 | 0.003 | 0.003 | 0.022 | 0.042 | 0.0018 |
| | **4-5** | 8E-15 | 2E-16 | 2E-16 | 2E-16 | 7E-13 | 2E-16 | 1E-04 | 2E-04 | 2E-06 | 3E-05 | 1E-04 | 5E-06 |
| **CbA** | **1-2** | - | - | - | - | - | - | - | - | - | - | - | - |
| | **1-3** | - | - | - | - | - | - | - | - | - | - | - | - |
| | **1-4** | - | - | - | - | - | - | 0.02 | - | - | - | - | - |
| | **1-5** | 2E-04 | 2E-04 | 9E-04 | 0.004 | 0.011 | 5E-04 | 2E-16 | 2E-16 | 3E-13 | 3E-10 | 5E-10 | 8E-16 |
| | **2-3** | - | - | - | - | - | - | - | - | - | - | - | - |
| | **2-4** | - | - | - | - | - | - | 0.028 | - | - | - | - | - |
| | **2-5** | 1E-05 | 4E-08 | 2E-07 | 1E-06 | 4E-06 | 7E-08 | 2E-16 | 2E-16 | 2E-16 | 8E-15 | 5E-15 | 2E-16 |
| | **3-4** | - | - | - | - | - | - | - | - | - | - | - | - |
| | **3-5** | 2E-05 | 2E-07 | 3E-06 | 1E-05 | 2E-05 | 6E-07 | 7E-16 | 2E-16 | 4E-14 | 3E-11 | 1E-11 | 2E-16 |
| | **4-5** | 0.009 | 5E-07 | 9E-07 | 3E-06 | 1E-05 | 3E-06 | 2E-11 | 2E-16 | 2E-16 | 1E-13 | 6E-14 | 2E-16 |
| **CbP** | **1-2** | - | - | - | - | - | - | - | - | - | - | - | - |
| | **1-3** | - | - | - | - | - | - | 0.008 | 0.036 | - | - | - | 0.0338 |
| | **1-4** | - | - | - | - | - | - | 0.029 | - | - | - | - | - |
| | **1-5** | 9E-08 | 7E-07 | 9E-06 | 2E-05 | 1E-04 | 3E-07 | 2E-16 | 2E-11 | 3E-08 | 8E-07 | 1E-05 | 3E-12 |
| | **2-3** | - | - | - | - | - | - | - | - | - | - | - | - |
| | **2-4** | - | - | - | - | - | - | - | - | - | - | - | - |
| | **2-5** | 2E-06 | 3E-06 | 6E-06 | 2E-05 | 6E-05 | 7E-07 | 2E-16 | 3E-10 | 3E-08 | 8E-07 | 1E-05 | 3E-11 |
| | **3-4** | - | - | - | - | - | - | - | - | - | - | - | - |
| | **3-5** | 1E-08 | 2E-07 | 2E-06 | 6E-06 | 2E-05 | 5E-08 | 3E-14 | 7E-07 | 7E-05 | 3E-04 | 0.002 | 1E-07 |
| | **4-5** | 1E-05 | 2E-06 | 3E-06 | 5E-06 | 2E-05 | 4E-07 | 6E-16 | 8E-10 | 6E-08 | 8E-07 | 9E-06 | 8E-11 |
| **AI** | **1-2** | - | 1E-04 | 4E-05 | 1E-05 | 2E-06 | 1E-04 | 0.004 | - | - | - | - | - |
| | **1-3** | - | 0.018 | 0.008 | 0.004 | 8E-04 | 0.012 | - | - | - | - | - | - |
| | **1-4** | - | 1E-04 | 2E-05 | 3E-06 | 4E-07 | 2E-05 | 0.042 | - | - | - | - | - |
| | **1-5** | - | - | - | 0.015 | 0.004 | 0.038 | 7E-06 | 2E-15 | 3E-16 | 2E-16 | 2E-16 | 2E-16 |
| | **2-3** | - | - | - | - | - | - | 0.01 | - | - | - | - | - |
| | **2-4** | - | - | - | - | - | - | - | - | - | - | - | - |
| | **2-5** | - | 0.018 | 0.014 | 0.031 | 0.046 | - | - | 3E-12 | 1E-15 | 2E-16 | 2E-16 | 1E-13 |
| | **3-4** | - | - | - | 0.048 | - | - | - | - | - | - | - | - |
| | **3-5** | - | - | - | - | - | - | 2E-05 | 3E-16 | 2E-16 | 2E-16 | 2E-16 | 2E-16 |
| | **4-5** | - | 0.016 | 0.007 | 0.015 | 0.018 | 0.015 | 0.008 | 2E-12 | 2E-15 | 2E-16 | 2E-16 | 8E-15 |
| **OB** | **1-2** | - | - | - | - | - | - | - | - | - | - | - | - |
| | **1-3** | - | - | - | - | - | - | 0.049 | - | - | - | - | - |
| | **1-4** | - | - | - | - | - | - | - | - | - | - | - | - |
| | **1-5** | 0.031 | 1E-04 | 5E-04 | 7E-05 | 8E-05 | 8E-05 | 0.02 | 1E-05 | 7E-05 | 8E-06 | 8E-06 | 9E-06 |
| | **2-3** | - | - | - | - | - | - | - | - | - | - | - | - |
| | **2-4** | - | - | - | - | - | - | - | - | - | - | - | - |
| | **2-5** | - | 1E-04 | 3E-04 | 2E-05 | 2E-05 | 5E-05 | 0.036 | 1E-05 | 3E-05 | 2E-06 | 1E-06 | 4E-06 |

| | | | | | | | | | | | | | |
|---|---|---|---|---|---|---|---|---|---|---|---|---|---|
| | **3-4** | - | - | - | - | - | - | - | - | - | - | - | - |
| | **3-5** | - | 0.001 | 0.004 | 6E-04 | 6E-04 | 0.004 | - | 2E-04 | 8E-04 | 9E-05 | 7E-05 | 0.0007 |
| | **4-5** | - | 3E-04 | 5E-04 | 5E-05 | 5E-05 | 1E-04 | 0.037 | 2E-05 | 4E-05 | 3E-06 | 2E-06 | 6E-06 |
| **Obpost** | **1-2** | - | - | - | - | - | - | - | - | - | - | - | - |
| | **1-3** | - | - | - | - | - | - | - | - | - | - | - | - |
| | **1-4** | 0.031 | - | - | - | - | - | 0.029 | - | - | - | - | - |
| | **1-5** | 3E-04 | 2E-08 | 4E-07 | 7E-09 | 2E-08 | 5E-09 | 1E-06 | 2E-16 | 2E-16 | 2E-16 | 2E-16 | 2E-16 |
| | **2-3** | - | - | - | - | - | - | - | - | - | - | - | - |
| | **2-4** | - | - | - | - | - | - | - | - | - | - | - | - |
| | **2-5** | 9E-04 | 2E-08 | 2E-07 | 1E-09 | 1E-09 | 2E-09 | 2E-06 | 2E-16 | 2E-16 | 2E-16 | 2E-16 | 2E-16 |
| | **3-4** | - | - | - | - | - | - | - | - | - | - | - | - |
| | **3-5** | 0.005 | 7E-09 | 6E-07 | 7E-09 | 9E-09 | 2E-08 | 4E-06 | 2E-16 | 2E-16 | 2E-16 | 2E-16 | 2E-16 |
| | **4-5** | - | 2E-07 | 3E-06 | 4E-08 | 1E-07 | 8E-07 | 0.004 | 2E-16 | 2E-16 | 2E-16 | 2E-16 | 2E-16 |
| **Obmed** | **1-2** | - | - | - | - | - | - | - | - | - | - | - | - |
| | **1-3** | - | - | - | - | - | - | - | - | - | - | - | - |
| | **1-4** | - | - | - | - | - | - | - | - | - | - | - | - |
| | **1-5** | 0.002 | 1E-05 | 1E-04 | 1E-05 | 2E-05 | 5E-06 | 9E-05 | 3E-12 | 3E-11 | 4E-13 | 4E-13 | 4E-13 |
| | **2-3** | - | - | - | - | - | - | - | - | - | - | - | - |
| | **2-4** | - | - | - | - | - | - | - | - | - | - | - | - |
| | **2-5** | 0.009 | 2E-05 | 8E-05 | 4E-06 | 4E-06 | 4E-06 | 5E-04 | 4E-12 | 6E-12 | 3E-14 | 2E-14 | 2E-13 |
| | **3-4** | - | - | - | - | - | - | - | - | - | - | - | - |
| | **3-5** | - | 2E-05 | 3E-04 | 3E-05 | 3E-05 | 5E-05 | 0.002 | 2E-12 | 4E-11 | 3E-13 | 2E-13 | 2E-12 |
| | **4-5** | 0.002 | 8E-06 | 5E-05 | 3E-06 | 3E-06 | 2E-06 | 5E-05 | 3E-13 | 2E-12 | 9E-15 | 8E-15 | 1E-14 |
| **OC** | **1-2** | 3E-05 | - | - | - | - | - | 1E-09 | 0.008 | - | - | - | 0.0118 |
| | **1-3** | - | - | - | - | - | - | - | - | - | - | - | - |
| | **1-4** | 2E-08 | 3E-04 | - | - | - | 0.021 | 3E-13 | 2E-04 | - | - | - | 0.0029 |
| | **1-5** | 5E-13 | 0.006 | 0.029 | - | - | 3E-04 | 2E-16 | 4E-16 | 6E-15 | 8E-13 | 2E-10 | 2E-16 |
| | **2-3** | 0.003 | - | - | - | - | - | 1E-08 | 0.015 | - | - | - | 0.0116 |
| | **2-4** | - | 0.02 | - | - | - | - | - | - | - | - | - | - |
| | **2-5** | 9E-04 | - | 0.037 | - | 0.031 | 0.007 | 8E-15 | 3E-09 | 8E-13 | 1E-11 | 1E-10 | 3E-15 |
| | **3-4** | 4E-06 | 0.005 | - | - | - | - | 2E-12 | 3E-04 | - | - | - | 0.0025 |
| | **3-5** | 2E-10 | - | - | - | - | 0.002 | 2E-16 | 2E-16 | 2E-16 | 2E-14 | 1E-12 | 2E-16 |
| | **4-5** | - | - | - | - | - | - | 1E-10 | 2E-06 | 5E-11 | 2E-11 | 1E-10 | 2E-13 |
| **midVermis** | **1-2** | - | - | - | - | - | - | | | | | | |
| | **1-3** | - | - | - | - | - | - | | | | | | |
| | **1-4** | 0.014 | - | - | - | - | - | | | | | | |
| | **1-5** | 2E-05 | 2E-04 | 6E-04 | 0.005 | 0.009 | 2E-04 | | | | | | |
| | **2-3** | - | - | - | - | - | - | | | | | | |
| | **2-4** | 0.002 | - | - | - | - | - | | | | | | |
| | **2-5** | 3E-07 | 8E-09 | 2E-08 | 6E-07 | 9E-07 | 6E-09 | | | | | | |
| | **3-4** | 1E-03 | - | - | - | - | - | | | | | | |
| | **3-5** | 1E-07 | 4E-08 | 3E-07 | 4E-06 | 5E-06 | 3E-08 | | | | | | |
| | **4-5** | 0.047 | 1E-07 | 2E-07 | 4E-06 | 8E-06 | 6E-06 | | | | | | |
| **midOB** | **1-2** | - | - | - | - | - | - | | | | | | |
| | **1-3** | - | - | - | - | - | - | | | | | | |
| | **1-4** | - | - | - | - | - | - | | | | | | |
| | **1-5** | 0.036 | 1E-04 | 5E-04 | 7E-05 | 8E-05 | 8E-05 | | | | | | |
| | **2-3** | - | - | - | - | - | - | | | | | | |
| | **2-4** | - | - | - | - | - | - | | | | | | |
| | **2-5** | - | 1E-04 | 2E-04 | 2E-05 | 2E-05 | 4E-05 | | | | | | |
| | **3-4** | - | - | - | - | - | - | | | | | | |
| | **3-5** | - | 0.001 | 0.004 | 6E-04 | 5E-04 | 0.003 | | | | | | |
| | **4-5** | - | 2E-04 | 3E-04 | 3E-05 | 3E-05 | 7E-05 | | | | | | |

### Supplementary Table 1.

### EXTENDED LEGENDS

**Supplementary Table 1. Simple effects contrasts between vaccine groups and each ROI.** Group 1, controls; group 2, 0.075 μg PsIV; group 3, 0.75 μg PsIV; group 4, 3.75 μg PsIV; group 5, prime boost. Entries are p-values, a '-' denotes insignificance. See Methods for ROI descriptions.
